# Multiomics analysis revealed anti-freezing mechanism of *Staphylococcus aureus* in its anti-freezing strain

**DOI:** 10.1101/2025.01.30.635745

**Authors:** Wu Youzhi, Sun Linjun, Shi Chunlei, Wu xianghuan, Jiao Lingxia, Ye Fuzhou

**Author notes:** Corresponding Author: Jiao Lingxia Contact Number.

## Abstract

Frozen food is currently a common food type. However, the presence of *Staphylococcus aureus* contamination caused serious challenge to frozen food safety. In this study, we explored the differences between sensitive strains and anti-freeze strains through multiomics analysis such as proteomics, phosphorylated proteomics, and metabolomics studies to understand the anti-freezing mechanism of *S. aureus*. This study compared the proteomics, phosphorylated proteomics and metabolomic differences between anti-freeze strains and sensitive strains before and after freezing. pre- and post-freezing, antifreeze strains mainly responded to low temperatures by enhancing antioxidant capacity, energy metabolism and antibiotic synthesis before freezing, while after freezing, bacteria depended more on biofilm formation, nitrogen metabolism and amino acid metabolism to improve frost resistance and adaptability. The *ket* gene such as *sucD* and *coaD* in the energy metabolism and metabolic pathways, *gapA1* and *tpiA* in glucose metabolism and energy balance, *asd* and *gnd* in the metabolic pathways of amino acid synthesis and protein synthesis, *crtN* in the context of protective mechanisms and stress response should be explored furthermore. In metabolomic research, the arginine synthesis pathway, biofilm formation pathway, Adenosine triphosphate-binding cassette transporters and other metabolic pathways deserve special attention and research.

**IMPORTANCE:** This study highlights the importance of understanding the anti-freezing mechanisms of *Staphylococcus aureus* in frozen food safety. By analyzing proteomics, phosphorylated proteomics, and metabolomics, it identifies key pathways like energy metabolism, biofilm formation, and amino acid synthesis that enhance bacterial frost resistance, offering insights for improving food preservation and safety.

## INTRDUCTION

The rapid economic growth and the quickening tempo of modern life have led to a shift in dietary habits, favoring simplicity and efficiency. Consequently, quick-frozen foods have emerged as a popular choice among consumers for their convenience.(1) However, the presence of *Staphylococcus aureus* in these products has become a significant concern for the frozen food industry. Data from the Jilin Provincial Center for Disease Control and Prevention reveals that between 2016 and 2019, *S. aureus* was detected in 8 out of 13 types of foods, with a particularly high detection rate of 9% in quick-frozen rice and flour products(2). European Union statistics indicate that at least 800 foodborne outbreaks in the 1990s were attributed to *S. aureus* (3). According to European Food Safety Authority, 40 out of 645 food samples from five countries tested positive for staphylococcal enterotoxins in 2017 (4). In the United States, *S. aureus* is responsible for approximately 240,000 cases of food poisoning annually (5). From 500 raw meat samples (chicken n=130, turkey n=130, fish n=240) collected in Akure from Sep 2020 to Dec 2021, 125 *S. aureus* isolates were identified, with the highest prevalence (40.80%) in turkey meat (6).

Furthermore, *S. aureus* can survive for up to 7 years under frozen conditions(7, 8). Studies have shown that the bacterium can endure at 10°C for at least 480 hours in sliced cooked chicken breast (7), and it remains viable at temperatures below -20° C, which are typical for frozen food storage (9). The production of enterotoxins by *S. aureus* during its survival in frozen foods can lead to prolonged contamination, with these toxins being the primary cause of staphylococcal food poisoning (10). The frequent isolation of *S. aureus* strains carrying enterotoxin genes from food samples underscores the gravity of the issue (11). A prevalence study in China found that 60% of quick-frozen dumplings were positive for *S. aureus*, with 10.3% to 38.5% of these strains harboring enterotoxin genes (12). However, at present, the antifreeze mechanism of *S. aureus* is still unclear, and further exploration is needed. This highlights the urgent need to understand the anti-freeze mechanisms of *S. aureus* in quick-frozen foods to inform food processing strategies aimed at preventing contamination.

In this study, we utilized anti-freezing and sensitive strains of *S. aureus* identified in our previous research and compared these strains to investigate their responses to freezing conditions using a multiomics approach, including proteomics, phosphoproteomics, and metabolomics. Our findings contribute to the development of a potential anti-freezing mechanism for *S. aureus*, which could guide future food safety measures to mitigate the risk of *S. aureus* contamination.

## RESULTS

### Proteomic Analysis of Sensitive and anti-freezing Strains Before and After Freezing

Proteomic analysis was conducted to identify differentially expressed proteins before and after freezing in sensitive and anti-freezing strains. GO enrichment analysis was performed based on the biological functions in which these differential proteins are involved. Consistent results were observed in both sensitive and anti-freezing strains before and after freezing, with a higher proportion of differentially expressed proteins in biological processes such as metabolic processes and cellular processes. In cellular component analysis, differentially expressed proteins were primarily associated with cell, cell part, cell membrane, and cell membrane part. In molecular function analysis, proteins with catalytic activity and binding activity were highly represented (Fig 1&2).

**Fig. 1.**
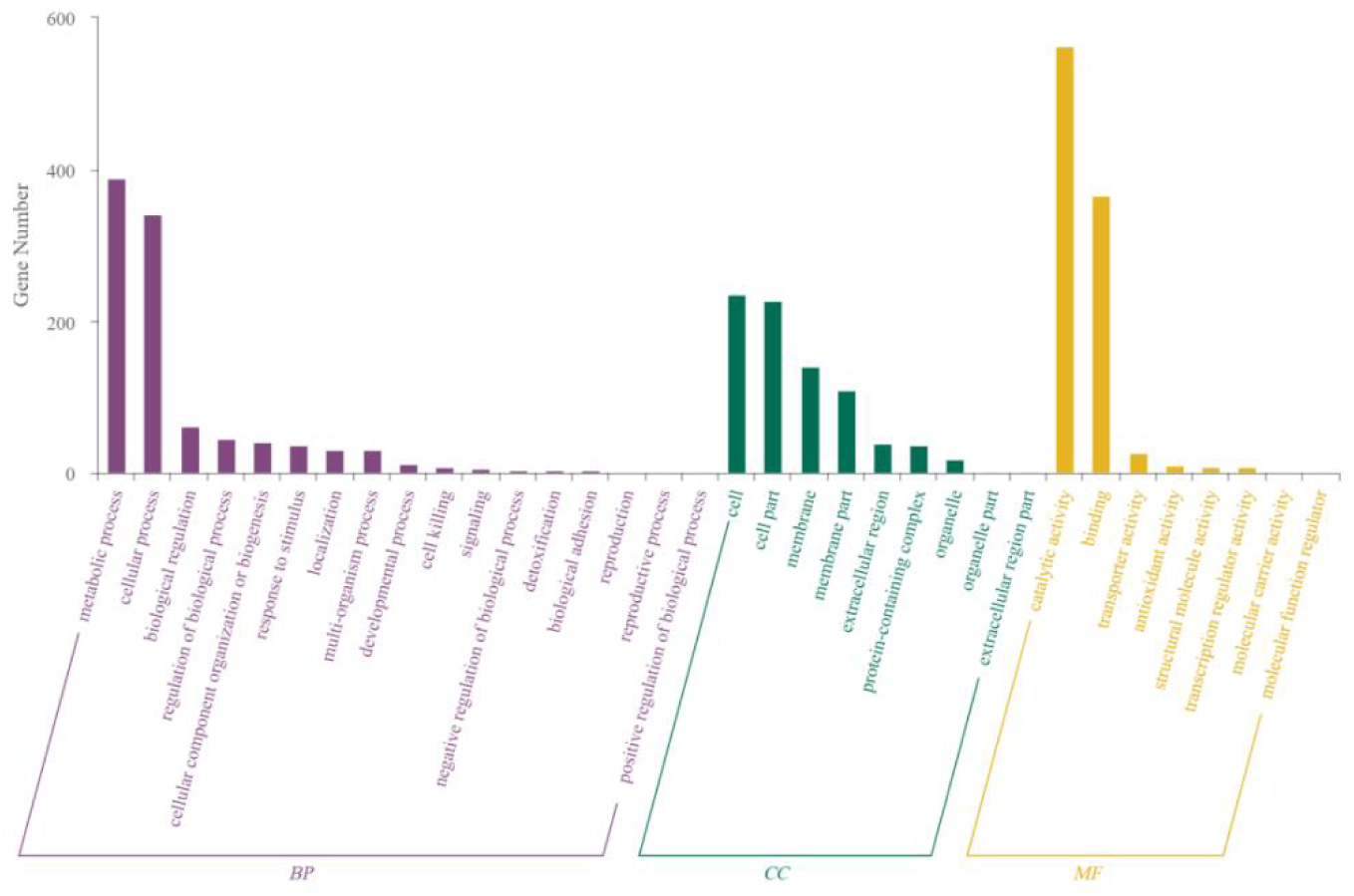
Statistical distribution of proteins corresponding to different modification sites for differential expressed proteins for sensitive and antifreeze strains before freezing in GO secondary classification before freezing through proteomic analysis.

**Fig. 2.**
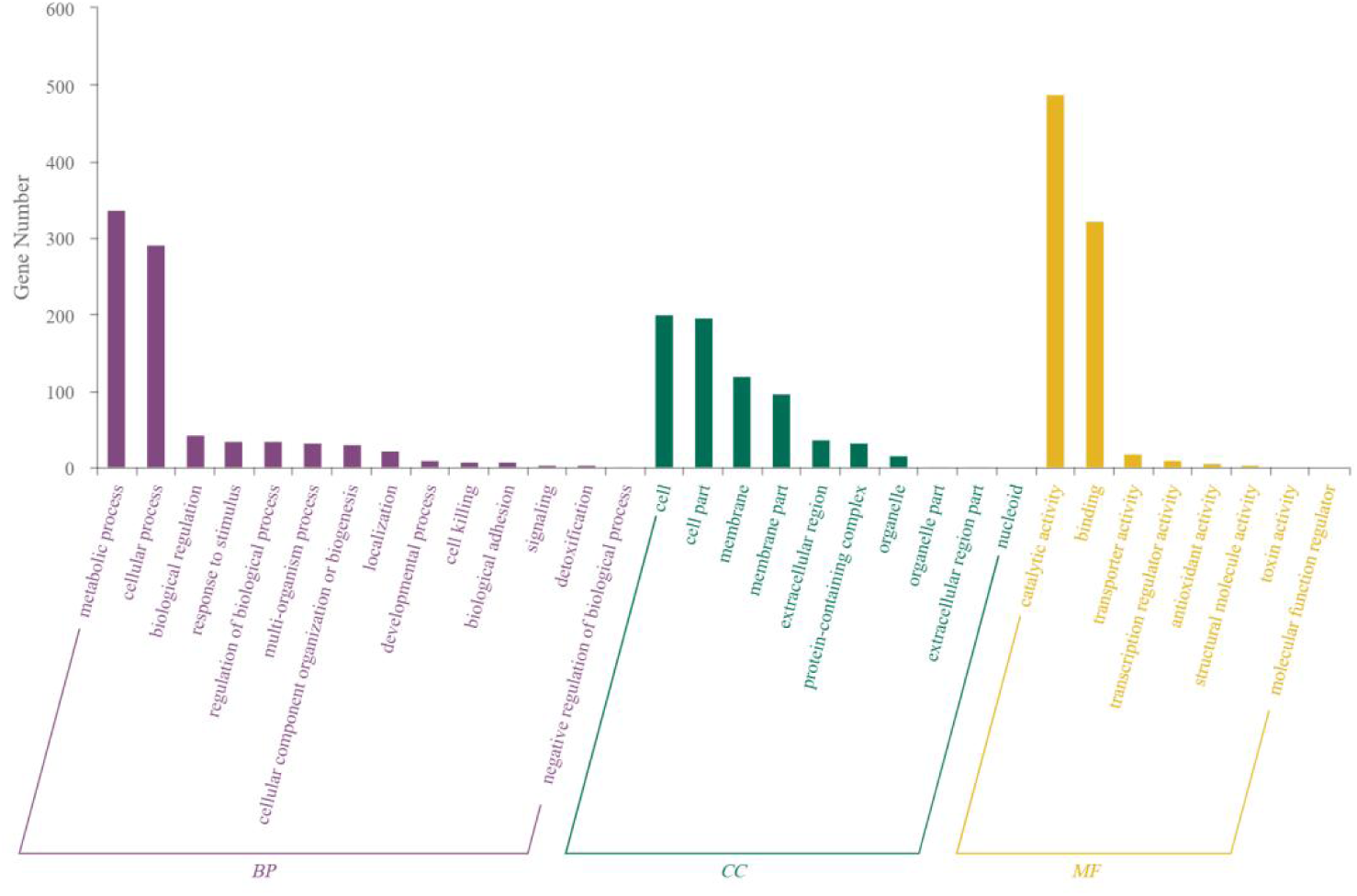
Statistical distribution of proteins corresponding to different modification sites for differential expressed proteins for sensitive and antifreeze strains before freezing in GO secondary classification after freezing through proteomic analysis

In this study, differential proteins related to sensitive and anti-freezing strains before freezing were annotated with KEGG to analyze the main metabolic and signal transduction pathways in which these proteins are involved. KEGG enrichment revealed that the top ten pathways for differentially expressed proteins in sensitive and anti-freezing strains before freezing included Novobiocin biosynthesis, Atrazine degradation, Arachidonic acid metabolism, Carotenoid biosynthesis, Other glycan degradation, Cyanoamino acid metabolism, Biosynthesis of ansamycins, Monobactam biosynthesis, C5-Branched dibasic acid metabolism, and Nicotinate and nicotinamide metabolism (Fig 3).

**Fig. 3.**
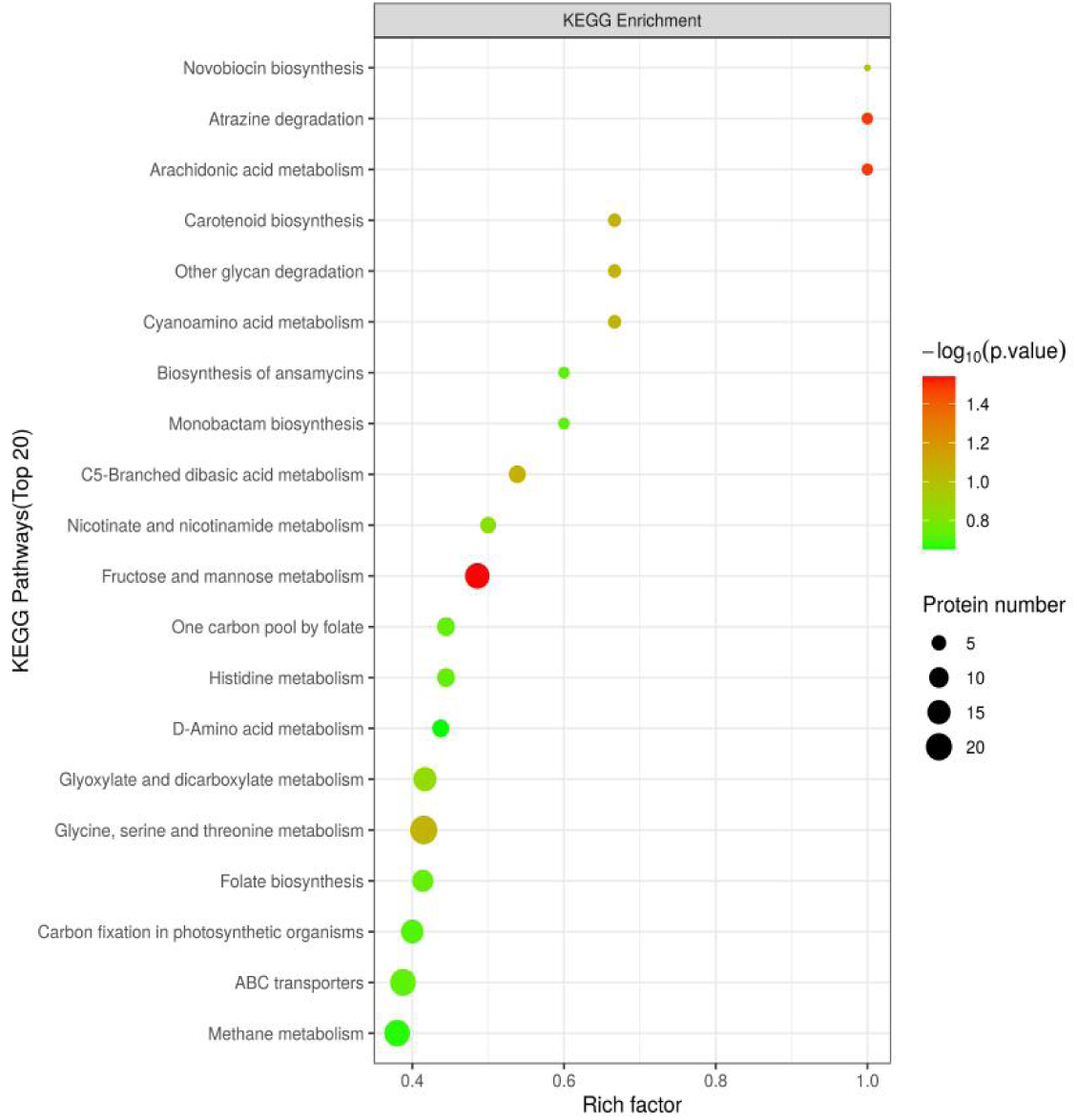
KEGG pathway annotation statistical chart of differentially expressed proteins between sensitive and antifreeze strains before freezing through proteomic analysis

After freezing, the top ten pathways for differentially expressed proteins in sensitive and anti-freezing strains changed to Atrazine degradation, Arachidonic acid metabolism, Streptomycin biosynthesis, Cyanoamino acid metabolism, C5-Branched dibasic acid metabolism, Biofilm formation - Vibrio cholerae, Cysteine and methionine metabolism, Nitrogen metabolism, Starch and sucrose metabolism, and Inositol phosphate metabolism (Fig 4).

**Fig. 4.**
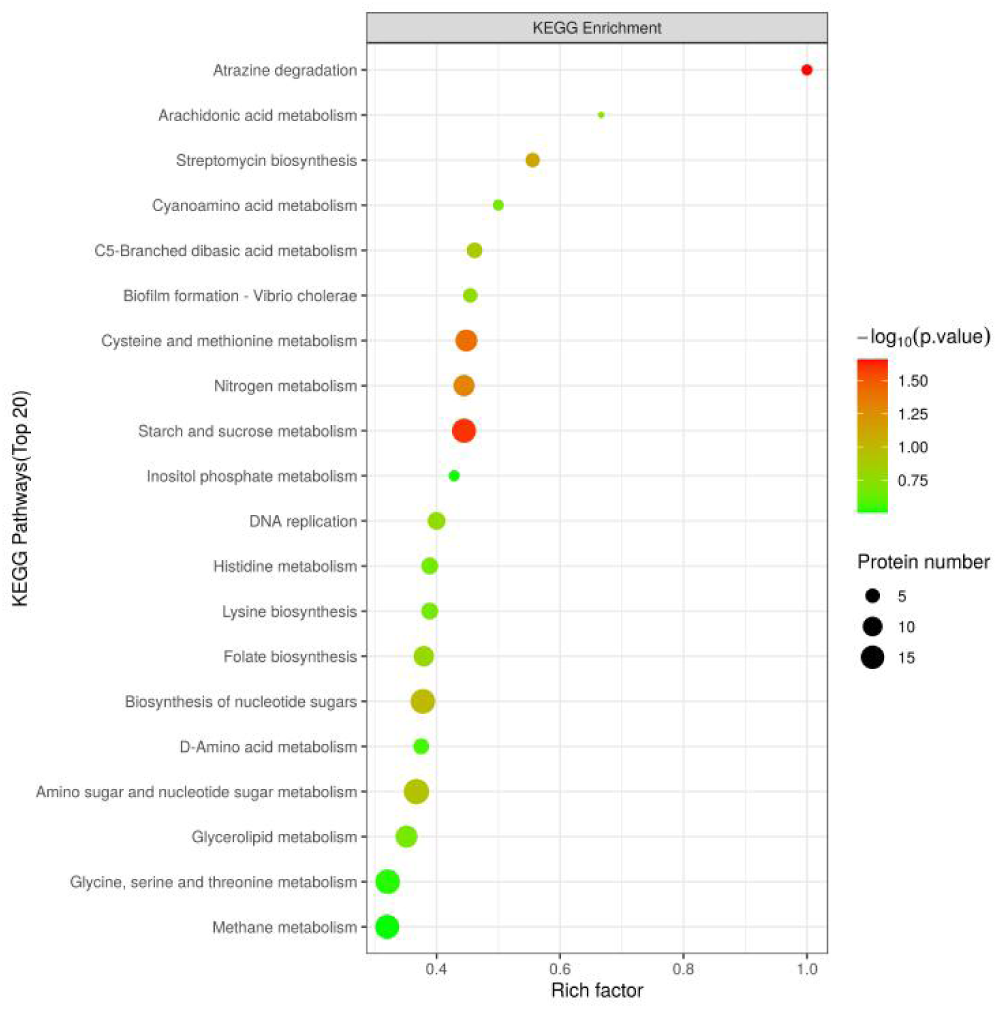
KEGG pathway annotation statistical chart of differentially expressed proteins between sensitive and antifreeze strains after freezing through proteomic analysis

Notably, the anti-freezing strain showed significant downregulation of *sucC* compared to the sensitive strain, while *sucD* was significantly upregulated. Additionally, before freezing, the anti-freezing strain’s *crtN*, *glyA*, and *tkt* were significantly downregulated compared to the sensitive strain, and after freezing, the *asd* gene in the anti-freezing strain was significantly upregulated compared to the sensitive strain.

### Phosphoproteomic Analysis of Sensitive and anti-freezing Strains Before and After Freezing

Phosphoproteomic analysis was conducted to identify differentially expressed proteins before and after freezing in sensitive and anti-freezing strains. GO enrichment analysis was performed based on the biological functions in which these differential proteins are involved. Consistent with proteomic results, a higher proportion of differentially expressed proteins were observed in cellular processes and metabolic processes. In cellular component analysis, differentially expressed proteins were mainly associated with cell part, cell, cell membrane part, and organelle. In molecular function analysis, proteins with catalytic activity and binding activity were highly represented (Fig 5 & Fig 6).

**Fig. 5.**
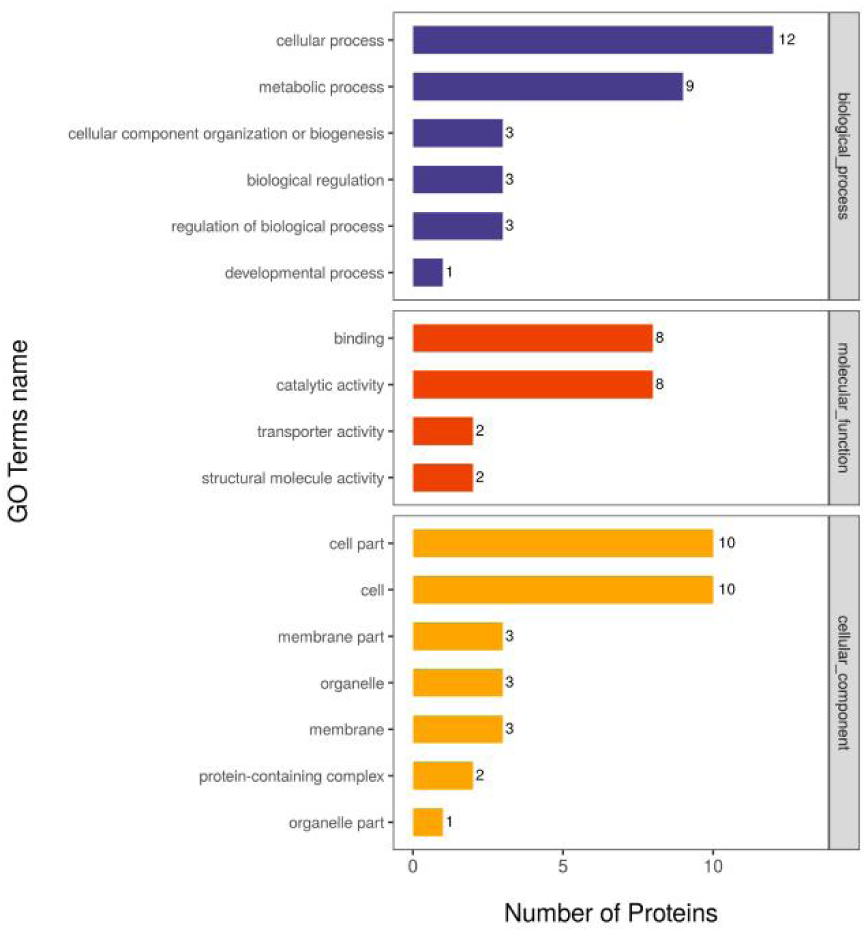
Statistical distribution of proteins corresponding to different modification sites for differential expressed proteins for sensitive and antifreeze strains before freezing in GO secondary classification through phosphoproteomic analysis

**Fig. 6.**
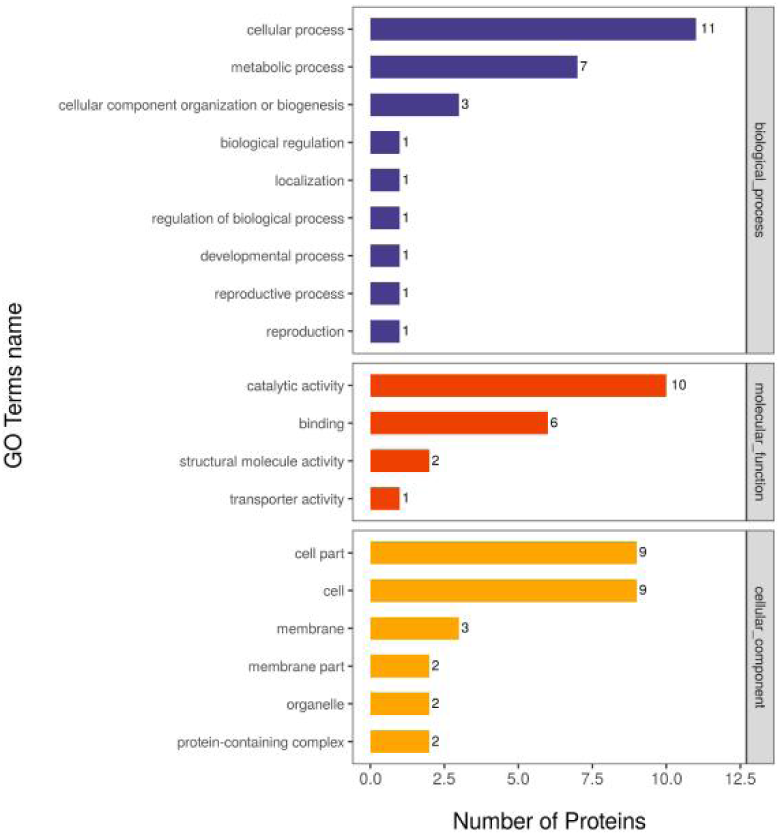
Statistical distribution of proteins corresponding to different modification sites for differential expressed proteins for sensitive and antifreeze strains after freezing in GO secondary classification through phosphoproteomic analysis

In this study, differential proteins related to sensitive and anti-freezing strains before freezing were annotated with KEGG to analyze the main metabolic and signal transduction pathways in which these proteins are involved. KEGG enrichment revealed that the top ten pathways for differentially expressed proteins in sensitive and anti-freezing strains before freezing included Pantothenate and CoA biosynthesis, Glycerolipid metabolism, Carbon fixation in photosynthetic organisms, HIF-1 signaling pathway, Fructose and mannose metabolism, Methane metabolism, Phosphotransferase system (PTS), Two-component system, Butanoate metabolism, and Glycerophospholipid metabolism (Fig 7).

**Fig. 7.**
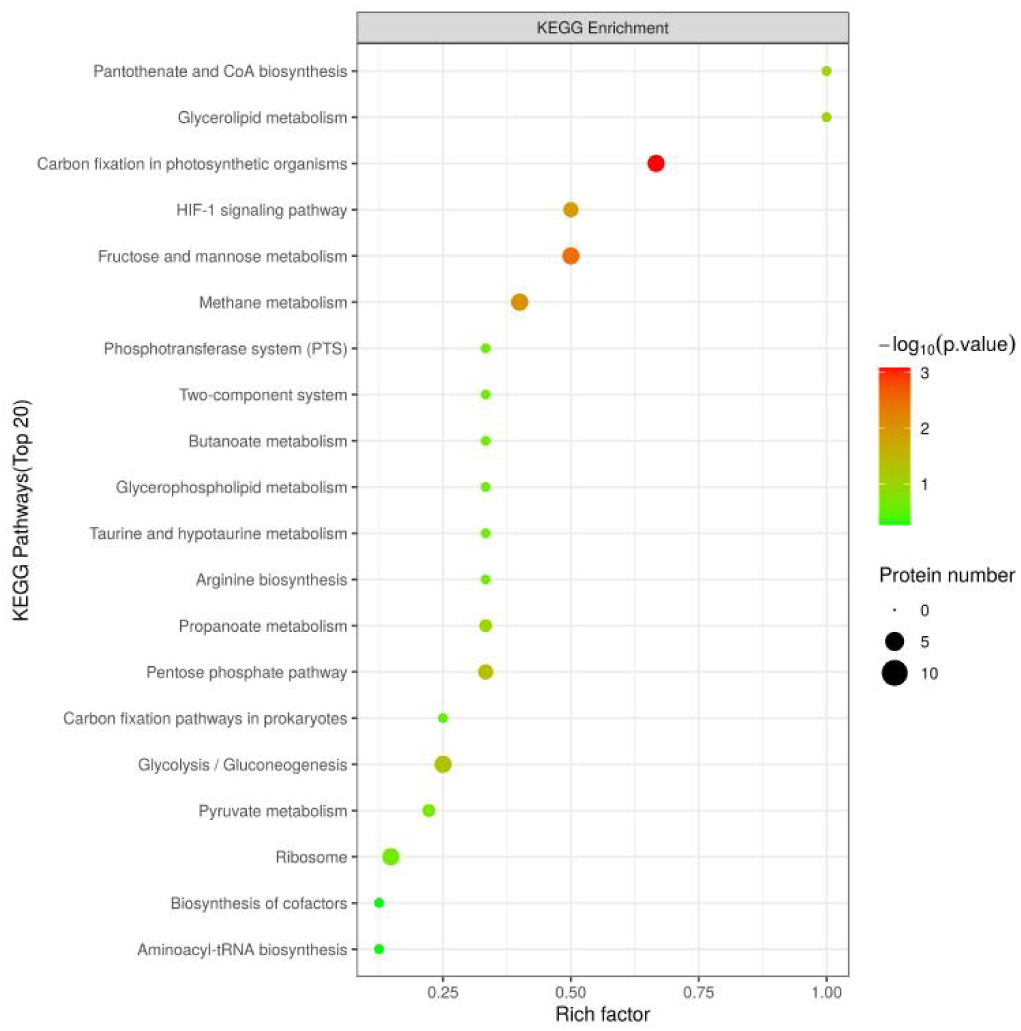
KEGG pathway annotation statistical chart of differentially expressed proteins between sensitive and antifreeze strains before freezing through phosphoproteomic analysis

After freezing, the top ten pathways for differentially expressed proteins in sensitive and anti-freezing strains changed to Nitrogen metabolism, Inositol phosphate metabolism, Glutathione metabolism, Carbon fixation in photosynthetic organisms, Fructose and mannose metabolism, Pentose phosphate pathway, Cell cycle - Caulobacter, Two-component system, Butanoate metabolism, and Taurine and hypotaurine metabolism (Fig 8).

**Fig. 8.**
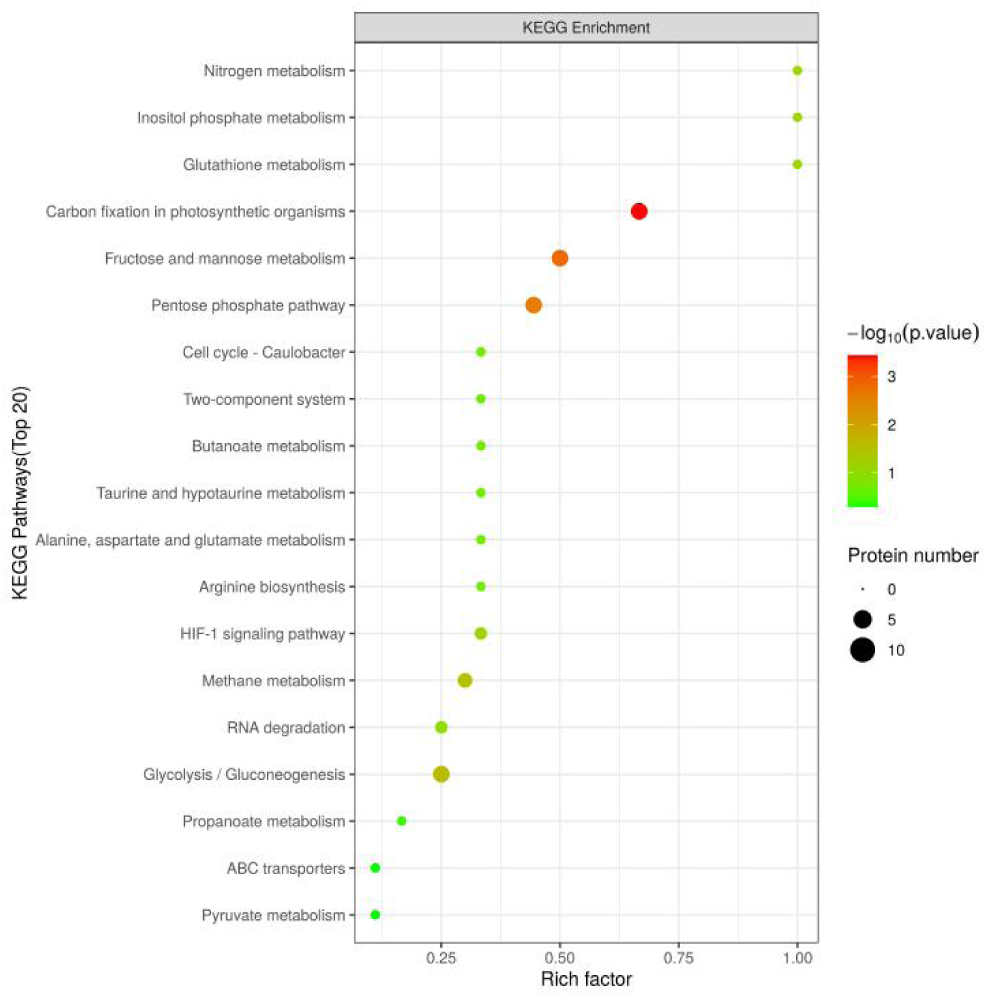
KEGG pathway annotation statistical chart of differentially expressed proteins between sensitive and antifreeze strains after freezing through phosphoproteomic analysis

Before freezing, *coaD* in the anti-freezing strain was significantly increased in Pantothenate and CoA biosynthesis. After freezing, *tpiA* in the anti-freezing strain was significantly upregulated in Inositol phosphate metabolism, Carbon fixation in photosynthetic organisms, and Fructose and mannose metabolism. *gnd* was significantly upregulated in Glutathione metabolism and Pentose phosphate pathway.

### Metabolomic Analysis of Sensitive and anti-freezing Strains Before and After Freezing

Initially, a VIP value greater than 1 was used as a screening criterion for differential metabolites between the two groups of samples. Subsequently, univariate statistical analysis was employed with a P-value less than 0.05 to verify the significance of differential metabolites. Metabolites with a VIP value greater than 1 and a P-value less than 0.05 were considered significantly differential metabolites, while those with a VIP value greater than 1 and a P-value less than 0.1 were considered differential metabolites. Results showed that before freezing, a total of 143 significantly differential metabolites were identified, with 92 metabolites significantly upregulated (Table S1) and 51 metabolites significantly downregulated (Table S2). After freezing, a total of 174 significantly differential metabolites were identified, with 110 metabolites significantly upregulated (Table S3) and 64 metabolites significantly downregulated (Table S4).

KEGG functional annotation revealed that before freezing, significantly differential metabolites were enriched in mTOR signaling pathway, GABAergic synapse, D-Arginine and D-Ornithine metabolism, Arginine biosynthesis, Protein digestion and absorption, Lysine degradation, cAMP signaling pathway, Alanine, aspartate and glutamate metabolism, and beta-Alanine metabolism (Fig 9). Although ABC transporters were not in the top ten, their metabolites also showed significant differences (*P*<0.05). After freezing, the pathways included mTOR signaling pathway, FoxO signaling pathway, GABAergic synapse, Protein digestion and absorption, Alanine, aspartate and glutamate metabolism, Arginine biosynthesis, D-Arginine and D-Ornithine metabolism, AMPK signaling pathway, and Cholinergic synapse (Fig 10).

**Fig. 9.**
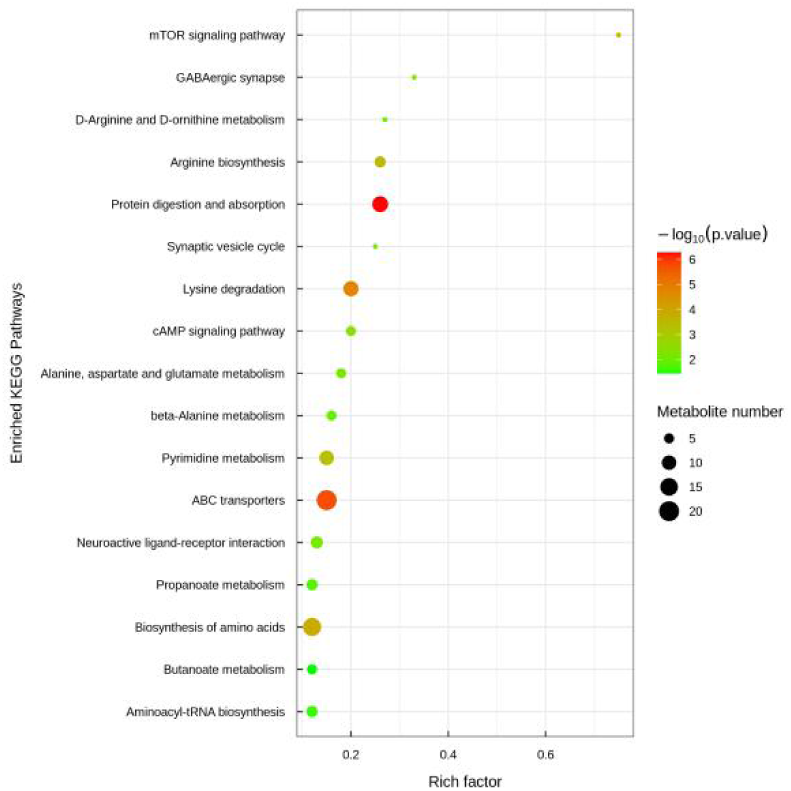
KEGG pathway annotation statistical chart of differentially expressed metabolites between sensitive and antifreeze strains before freezing through metabolites analysis

**Fig. 10.**
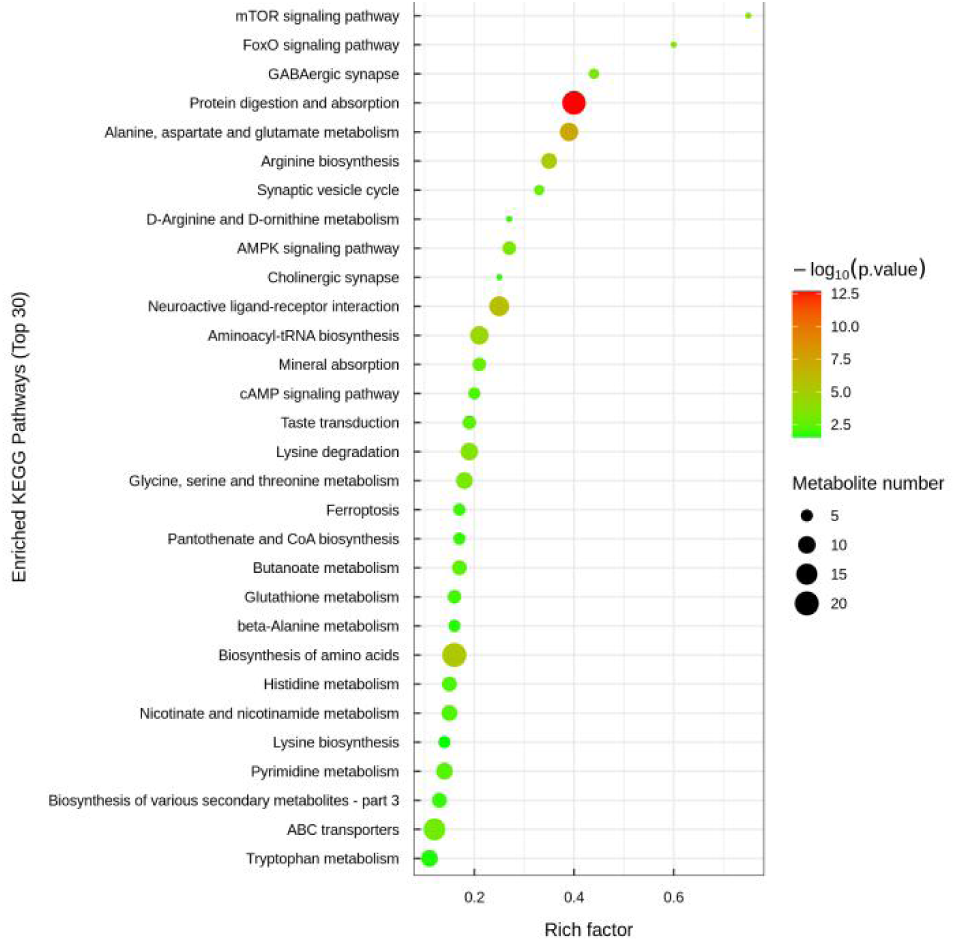
KEGG pathway annotation statistical chart of differentially expressed metabolites between sensitive and antifreeze strains after freezing through metabolites analysis

## DISCUSSION

*S. aureus* is a pathogenic bacterium that can survive in a variety of environments and cause infections. Not only can it adapt to room temperature, but it is also able to survive in extreme environments, including low-temperature conditions (e.g., freezing or cold) (13). Studying the antifreeze mechanism of *S. aureus* at low temperature is the key to prevent and control the survival and reproduction of *S. aureus* in quick-frozen foods. In this study, it was found that the frost resistance of *S. aureus* may be related to the stability of cell membranes, amino acid metabolism and protein synthesis, and the continuation of energy metabolism. In proteomics, it can be seen from the differential pathways from pre-freezing to post-freezing that the regulatory mechanism of metabolic pathways of *S. aureus* changes when faced with low temperatures, especially in energy metabolism, antioxidant response, amino acid metabolism and biofilm formation. Before freezing, antifreeze strains mainly responded to low temperatures by enhancing antioxidant capacity, energy metabolism and antibiotic synthesis, while after freezing, bacteria relied more on biofilm formation, nitrogen metabolism and amino acid metabolism to improve frost resistance and adaptability.

In the energy metabolism and metabolic pathways, the *sucD* gene in the antifreeze strain was significantly enhanced compared with the susceptible strain. The *sucD* gene is a gene associated with succinic acid metabolism and encodes a component of the succinate dehydrogenase complex. The Succinate Dehydrogenase Complex (SDH) is a key enzyme in the tricarboxylic acid cycle (TCA cycle) that catalyzes the oxidation of succinic acid to fumarate and transfers electrons to ubiquinone to produce ubiquinol. This process plays an important role in the energy metabolism of bacteria (14). At low temperatures, the rate of metabolism of bacteria usually decreases, but the activity of the succinate dehydrogenase complex is essential to maintain an adequate energy supply. *sucD* is higher in frost-resistant strains, which helps *S. aureus* to maintain the activity of the succinic acid cycle at low temperatures, thus promoting a continuous supply of energy. The *coaD* gene encodes an enzyme in coenzyme A synthesis. Coenzyme A is the core molecule of a variety of metabolic reactions, and is involved in important physiological processes such as fatty acid metabolism and energy synthesis. Several studies based on plant and animal survival at low temperatures have shown that *coaD* has the property of helping cells adjust the composition of membrane fatty acids and maintain the fluidity and stability of membranes (15–17). It is inferred that the *coaD* gene may also contribute to the survival of *S. aureus* at low temperatures.

In glucose metabolism and energy balance, *gapA1* and *tpiA* genes were significantly up-regulated in antifreeze strains compared with susceptible strains. They encode glyceraldehyde-3-phosphate dehydrogenase (GAPDH) and triphosphate isomerase (TPI), respectively. *gapA1* encodes glyceraldehyde-3-phosphate dehydrogenase (GAPDH), a key enzyme in the glycolysis process that catalyzes the formation of glyceraldehyde 3-phosphoglyceraldehyde to 1,3-bisphoglycerate while transferring energy to high-energy phosphobonds. *tpiA* encodes triphosphate isomerase (TPI), another key enzyme in the glycolysis process that catalyzes the conversion of dihydroxyacetone phosphate to glyceraldehyde-3-phosphate. In a cryogenic environment, the activity of GAPDH helps bacteria maintain energy metabolism and supports basic life activities at low temperatures (18).

In the metabolic pathways of amino acid synthesis and protein synthesis, the *asd* gene is associated with lysine biosynthesis. Yang et al. (19) found that under the influence of Litsea cubeba essential oil, genes related to lysine synthesis in *S. aureus* were upregulated, including the *asd* gene. Lysine synthesis is also related to cell membrane and cell wall stress. It can be inferred that the upregulation of *asd* may protect *S. aureus* from damage under extreme conditions, such as low temperatures. The *gnd* gene, which encodes glucose-6-phosphate dehydrogenase (G6PD) in the pentose phosphate pathway, is also upregulated. This enzyme catalyzes important reactions that not only aid in sugar metabolism but also generate reducing equivalents of NADPH, maintaining the antioxidant capacity within cells (20, 21). The *gnd* gene is an acid-shock gene in *S. aureus* (22). However, cross-protection, such as acid pH heat, as well as homologous resistance, has been reported to occur (23). Most current research is based on cross-protection studies under high temperatures (23, 24). Therefore, it can be inferred that acid-shock genes may also provide cross-protection under low-temperature conditions.

In the context of protective mechanisms and stress response, the downregulation of the *crtN* gene is noteworthy. The *crtN* gene is associated with the synthesis of carotenoids, which are natural antioxidants that help cells cope with oxidative stress (25). Inhibition of *crtN* function can block the synthesis of staphylococcal pigments, thereby reducing the virulence of *S. aureus* (26). It can be inferred that the reduced expression of *crtN* in cold-resistant strains compared to sensitive strains may be due to the need for these strains to survive under low-temperature conditions, thereby reducing their virulence, leading to the downregulation of the *crtN* gene. However, the specific relationship between the downregulation of *crtN* before freezing and cold resistance still requires further experimental validation. Future research should further explore the expression patterns of *crtN* before and after freezing, as well as in different strains, and combine these findings with cellular physiological changes before and after freezing to reveal the molecular mechanisms of cold resistance in different strains. Additionally, gene knockout or overexpression experiments should be utilized to explore the specific role of *crtN* in cold resistance, which will provide deeper insights into cold tolerance.

From a metabolomics perspective, metabolic pathways involving protein metabolism and membrane transport are also points of interest in cold resistance, consistent with results from proteomics and phosphoproteomics. Metabolites such as N-acetylglutamate and guanidine were significantly upregulated, while arginine, N-alpha-acetylornithine, and glutamine were significantly differences in the arginine synthesis pathway. Glutamate and glutamine can interconvert, and glutamate can be converted to ornithine through a series of metabolic processes, entering the urea cycle. Meanwhile, ornithine can be converted to guanidine in the presence of ornithine aminotransferase. Based on the results, it can be inferred that the arginine synthesis pathway is active, leading to a significant increase in guanidine. Subsequently, guanidine can be converted to arginine via pathways involving alanine, aspartate, and glutamate metabolism, and then arginine can be cleaved into citrulline and ornithine by arginase. Citrulline can enter the important tricarboxylic acid cycle for metabolism. However, the metabolism of arginine exhibits an inhibitory state.

Research has shown that arginine metabolism has a significant impact on the formation of biofilms by *S. aureus*. During biofilm formation, *S. aureus* selectively acquires arginine from the external environment (27, 28). Furthermore, arginine is involved in various important physiological processes, including energy metabolism, protein synthesis, and stress regulation in *S. aureus* (29). In the arginine deiminase (ADI) pathway, arginine is the precursor of citrulline and can also generate ornithine. Therefore, acid-adapted *Lactobacillus delbrueckii* subsp. bulgaricus can survive under high salt (180 g/L NaCl) conditions (30). This suggests that arginine allows bacteria to survive in stressful environments. It is speculated that the generation of arginine may be related to the freeze resistance mechanism of *S. aureus*, but how it affects the freeze resistance mechanism is still unclear.

Additionally, arginine can also be metabolized through the pathways involving arginine and proline metabolism. In proteomics studies, it has been observed that cold-resistant strains exhibit significant differences in the biofilm formation pathway after freezing. This suggests that biofilm formation may be associated with the cold resistance of *S. aureus*.Studies have shown that increased levels of proline can “awaken” bacteria in a dormant state (31). In other words, under stressful conditions, proline can reduce the sensitivity of *S. aureus* to the external environment, thereby promoting the formation of freeze-resistant strains. In summary, significant changes were observed in various amino acids and related metabolites in the arginine metabolism pathway, indicating the important role of this pathway in the freeze resistance mechanism of *S. aureus*, which warrants further investigation.

Adenosine triphosphate-binding cassette transporters (ABC transporters) are membrane-bound proteins containing an ATP-binding cassette. They utilize the energy released from ATP hydrolysis to facilitate the transmembrane transport of substances from the external environment into the cell (32–35). ABC transporters play a crucial role in maintaining cell integrity and are important in cell differentiation, signal transduction, and pathogenesis(35). Lin Jieting et al. (36) found that the glycine betaine ABC transporter system in Halo bacillus is closely related to its salt adaptation under high salt stress conditions. This indicates the crucial role of ABC transporter systems in bacterial response to adverse environments.

*S. aureus* exhibits increased sensitivity to the external environment under freezing conditions. Compared to sensitive strains, freeze-resistant strains of *S. aureus* are enriched in ABC transporter systems, primarily involving differences in amino acid metabolism. Amino acids, as important nitrogen sources in bacterial growth metabolism, also participate in energy metabolism as carbon sources. The amino acid transport proteins in ABC transporter systems promote the absorption of various amino acids by bacteria to meet the metabolic requirements for growth and virulence (35). The changes in multiple metabolites in freeze-resistant strains of *S. aureus* can trigger alterations in ABC transporter systems, potentially leading to changes in the transport of multiple amino acids across the cell membrane, which significantly affect bacterial growth and virulence expression in freeze-resistant strains.

Secondary metabolites are diverse and often have antagonistic effects on other microorganisms (37). They can also regulate gene expression (38) and influence bacterial virulence(28, 39). Pathways related to secondary metabolite synthesis are highly susceptible to external influences. Metabolomics analysis in this study revealed changes in differential metabolites involved in secondary metabolite synthesis and metabolism pathways in freeze-resistant strains of *S. aureus*, although the connections between them have yet to be fully understood.

In conclusion, antifreeze strains mainly responded to low temperatures by enhancing antioxidant capacity, energy metabolism and antibiotic synthesis before freezing, while after freezing, bacteria depended more on biofilm formation, nitrogen metabolism and amino acid metabolism to improve frost resistance and adaptability.The *ket* gene such as *sucD* and *coaD* in the energy metabolism and metabolic pathways, *gapA1* and *tpiA* in glucose metabolism and energy balance, *asd* and *gnd* in the metabolic pathways of amino acid synthesis and protein synthesis, *crtN* in the context of protective mechanisms and stress response should be explored furthermore. In metabolomic research, the arginine synthesis pathway, biofilm formation pathway, Adenosine triphosphate-binding cassette transporters and other metabolic pathways deserve special attention and research.

## MATERIALS AND METHODS

### Bacterial Strains and Cultivation

Sensitive (No.7) and anti-freezing (No.8) *S. aureus* strains were cultured in TSB and TSA media at 37°C for 24 hours, then cryopreserved at -18°C. Groups were designated as pre-freezing (H7, H8) and post-freezing (Q7, Q8) with ten replicates each.

### Protein extraction and digestion

The sample was extracted by SDT buffer (4% (w/v) SDS, 100 mM Tris-HCl pH7.6, 0.1 M DTT) using lysis method to extract protein. The protein was then quantified by BCA method. The peptides were desalted with C18 cartridge. After the peptides were lyophilized, 40 μL of 0.1% formic acid solution was added to redissolve them, and the peptides were quantified (OD_280_).

### Enrichment of phosphorylated peptides

Each peptide solution was lyophilized in vacuum, and then enriched with the High-SelectTM Fe-NTA Phosphopeptides Enrichment Kit (Thermos Scientific). The enriched phosphorylated peptides were concentrated in vacuo and reconstituted with 20 μL 0.1% formic acid solution for mass spectrometry experiment.

### LC-MS/MS data acquisition

Each sample was separated using nanoelute HPLC liquid phase system with nanoliter flow rate. buffer solution A is 0.1% formic acid aqueous solution, and buffer B is 0.1% formic acid acetonitrile aqueous solution. The sample was separated through an analytical column (homemade column, 25 cm, ID75 μm, 1.9 μm, C18) with a flow rate of 300 nL/min and a column temperature of 50 °C.

After chromatographic separation, the samples were analyzed by mass spectrometry with a timsTOF Pro mass spectrometer. The detection method is using positive ion. The ion source voltage is set to 1.5 kV. MS and MS/MS are both detected and analyzed by TOF. The mass spectrometer scan range was set at 100-1700m/z. The data acquisition mode adopts the parallel accumulation serial fragmentation (PASEF) mode. After a mass spectrometer is collected, 10 times of PASEF mode is used to collect precursor ions. The cycle window time is 1.17 seconds, and the charge number is in the range of 0-5. The spectrum, dynamic exclusion time of the tandem mass spectrometry scan was set to 24 seconds to avoid repeated scans of precursor ions.

### Protein identification and quantitative analysis

The original data of mass spectrometry analysis was a RAW file, which was processed by MaxQuant software (version 1.5.3.17).

### Bioinformatics Analysis

GO and KEGG annotations were performed using Blast2GO and KAAS, respectively, with Fisher’s exact test for enrichment analysis. Metabolomics samples were collected from bacterial pellets post-cultivation, preprocessed, and analyzed using UHPLC-MS/MS. Data were processed with XCMS and analyzed for differential metabolites using R software.

### Collection of metabolomics testing samples

*S. aureus* sensitive strains (Strain 7) and anti-freezing strains (Strain 8), cultivated until the stationary phase, were separately inoculated onto 16-20 TSA-YE agar plates. Half of the plates were incubated overnight at 30°C in a CO_2_ incubator, while the other half were pre-frozen at -30°C to -35°C for 45 minutes and then transferred to - 18°C overnight. Approximately 400 μL of pre-chilled PBS buffer was aspirated onto each plate, and the colonies of the control strains on the TSA-YE agar plates were gently and repeatedly blown using a sterile pipette tip. The liquid from the TSA-YE agar plates was then transferred to clean 1.5 mL eppendorf tubes. The tubes were centrifuged at 12,000 rpm and 4°C for 3 minutes to collect the bacterial pellets. After discarding the supernatant, the bacterial pellets were washed at least three times with pre-chilled PBS buffer. Subsequently, the pellets were snap-frozen in liquid nitrogen for 5-10 minutes and immediately stored at -80°C in a freezer.

### Sample preprocessing

The entire thawing process should be carried out at 4°C. Approximately 1 mL of pre-chilled methanol/acetonitrile/water solution (2:2:2 v/v) was aspirated and mixed thoroughly. The mixture was sonicated in an ice-water bath for approximately 30 minutes. It was then left to stand in a -20°C freezer for about 10 minutes, followed by centrifugation at 14,000 g and 4°C for 20 mins. During the drying process of the supernatant, a vacuum condition should be provided. For the actual mass spectrometry analysis, the sample was reconstituted in 100 μL of acetonitrile-water solution (acetonitrile:water = 1:1 v/v). After vigorous mixing and centrifugation at 14,000g and 4°C for 15 mins, the supernatant was used for injection during the analysis.

### Chromatography-mass spectrometry analysis

After separation using the Vanquish LC Ultra-High Performance Liquid Chromatography (UHPLC) system, the samples were analyzed using the Q Exactive series mass spectrometer (Thermo fisher Scientific) in both positive and negative ion modes using electrospray ionization (ESI). The ESI source and mass spectrometry settings were as follows: auxiliary gas heater 1 (Gas 1): 60, auxiliary gas heater 2 (Gas 2): 60, curtain gas (CUR): 30 psi, ion source temperature: 600 °C, spray voltage (ISVF): ±5500 V (positive and negative modes); the first mass-to-charge ratio detection range: 80-1200 Da, resolution: 60,000, scan accumulation time: 100 ms. The second stage employed a data-dependent acquisition method, with a scanning range of 70-1200 Da, a resolution of 30,000, scan accumulation time: 50 ms, and a dynamic exclusion time of 4 s.

### Data Processing and Statistical Analysis

Experiments with anti-freezing and sensitive strains of *S. aureus* were conducted with three replicates. Paired t-tests in SPSS (version 11.0) were used to analyze the significance of differences between treatment and control groups, with significance levels set at *P*<0.05.

## ACKNOWLEDGEMENTS

This work was supported by Program for Innovative Research Team (in Science and Technology) in University of Henan Province,(Grant number 21IRTSTHN024, awarded to LJ), the Chenguang Program of Shanghai Education Development Foundation and Shanghai Municipal Education Commission(Grant number 22CGB08, awarded to XW) and National Natural Science Foundation of China, General Project, (Grant number 31972169, awarded to LC)

---

**Table S1.**
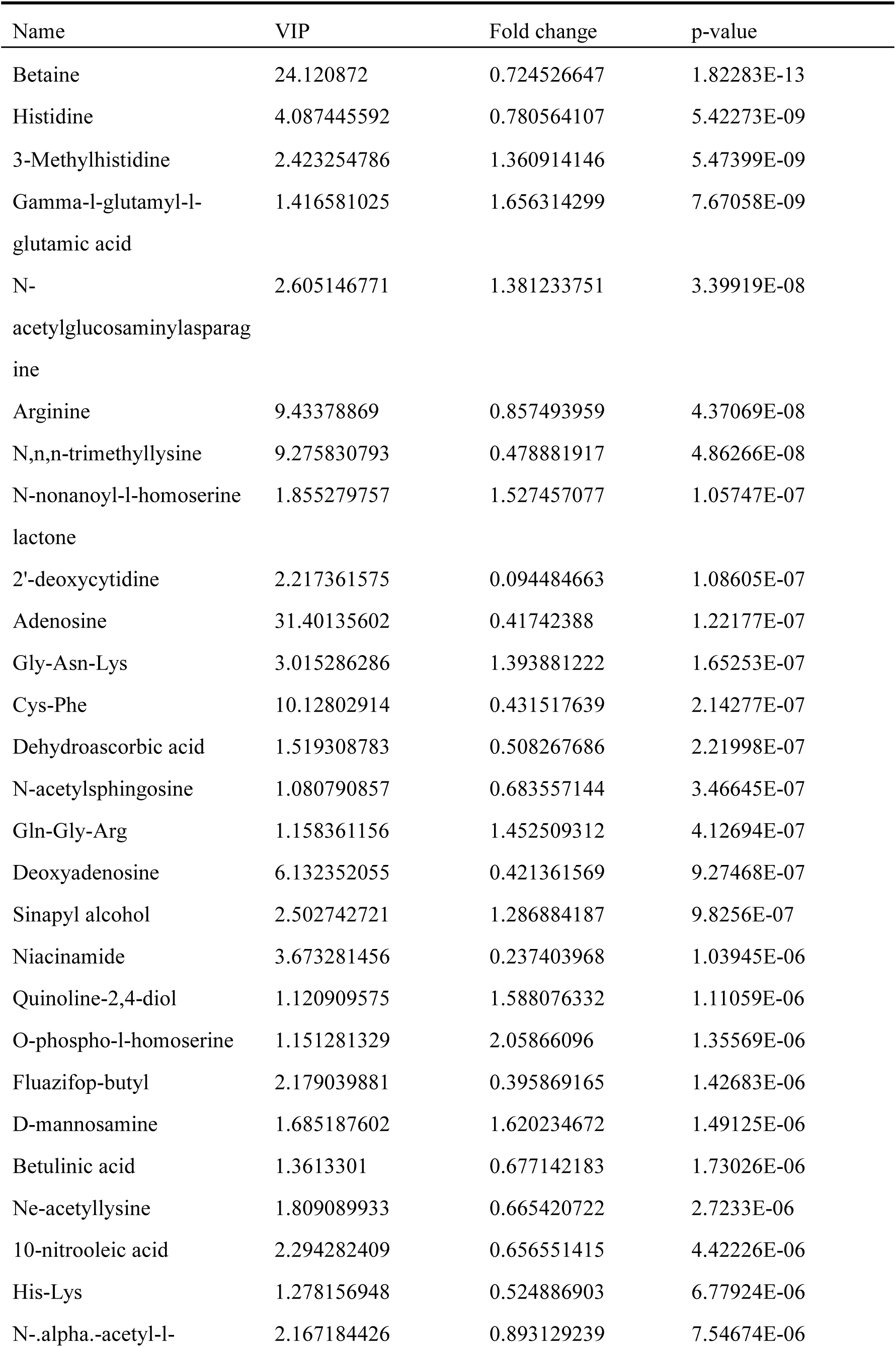
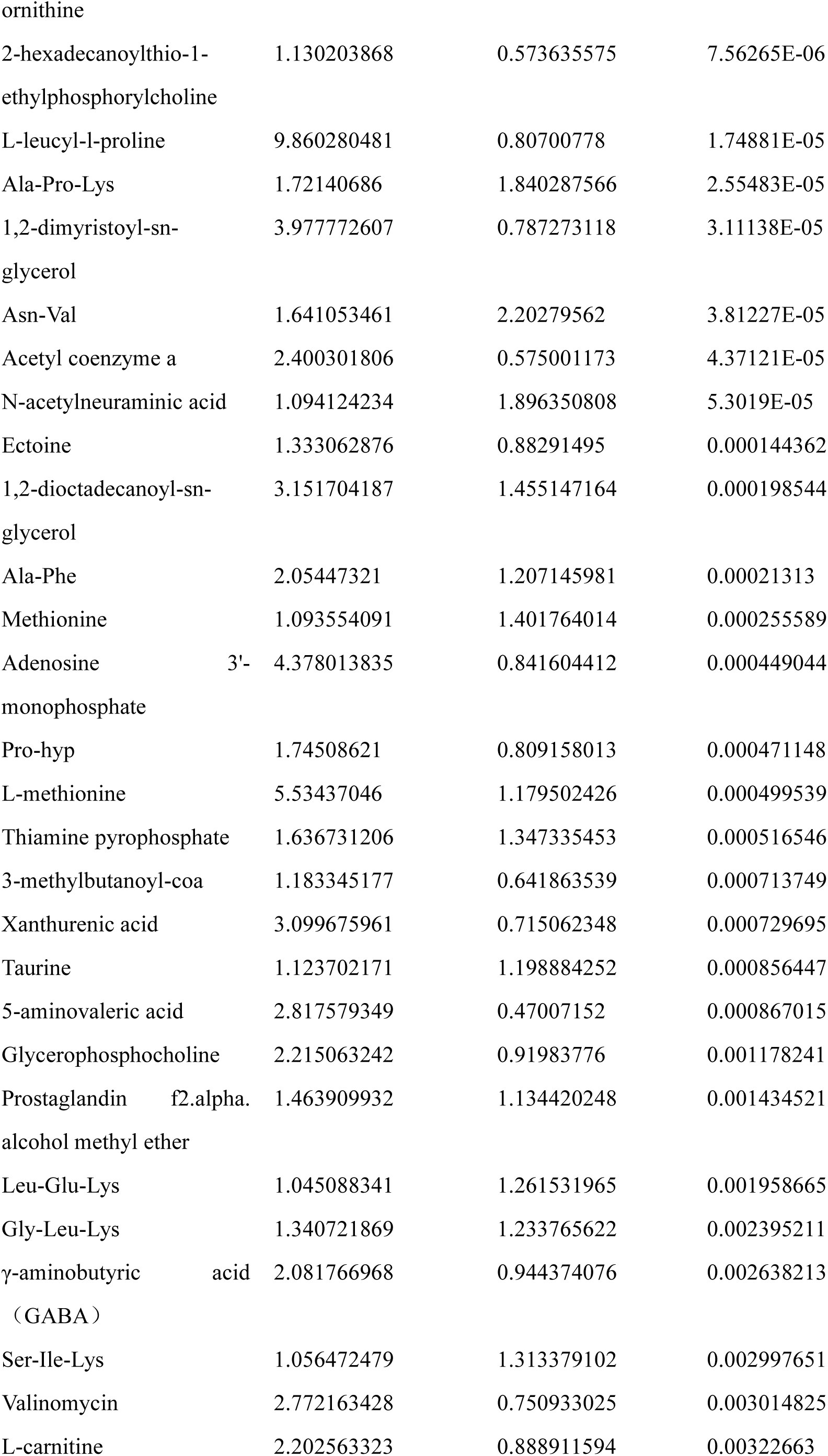
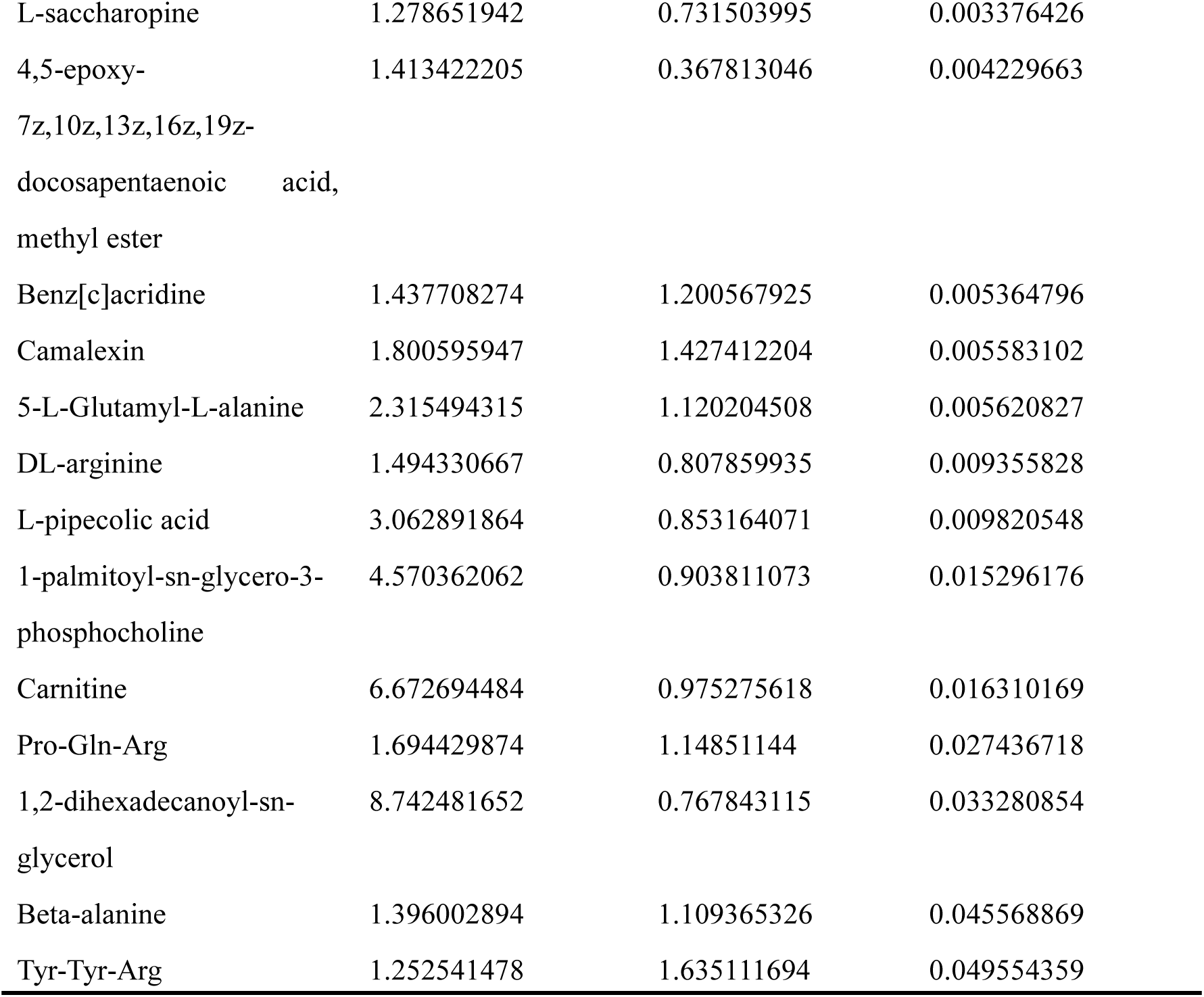
Staphylococcus aureus H7 vs H8 differential metabolites before freezing in positive ion mode.

**Table S2.**
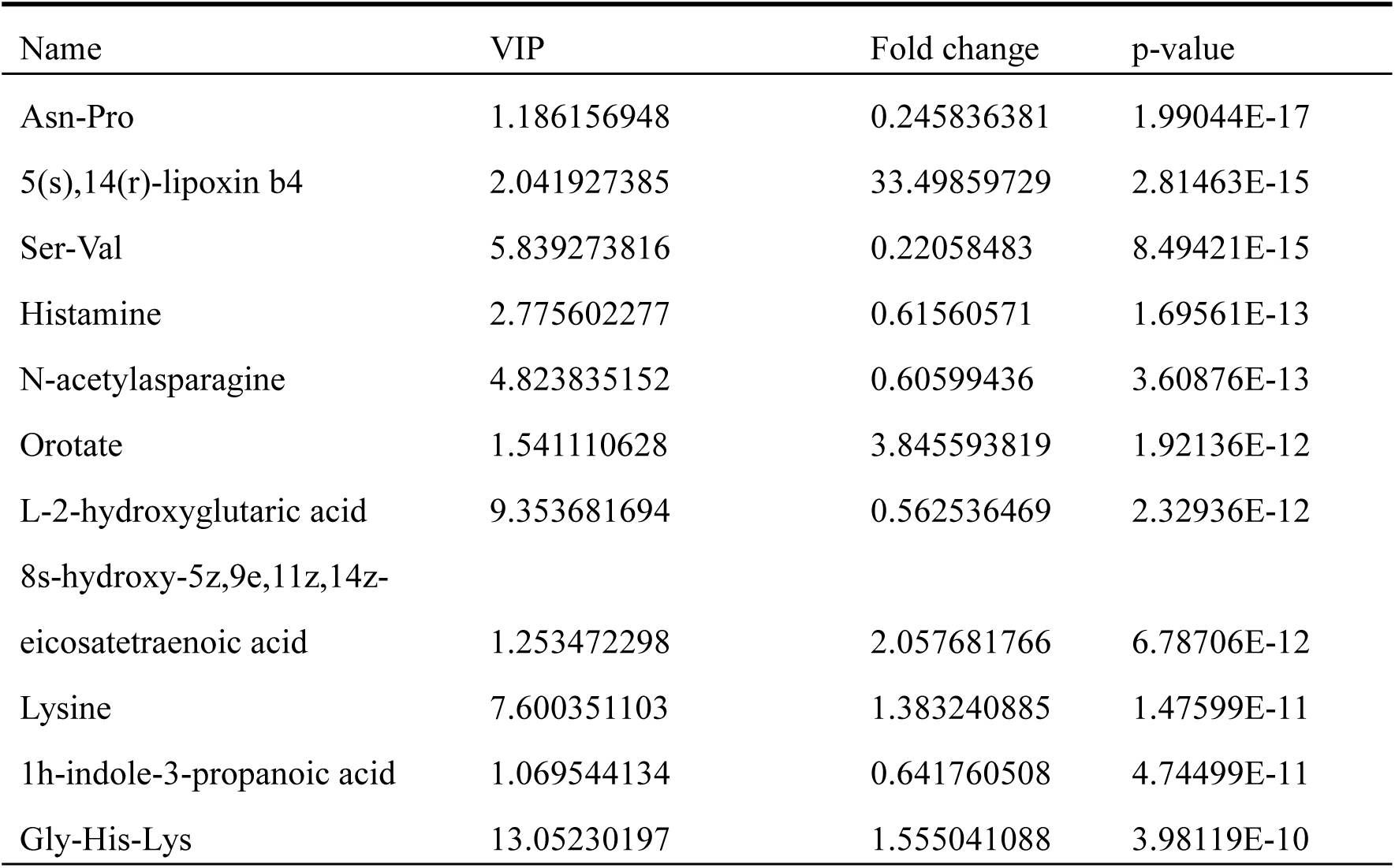
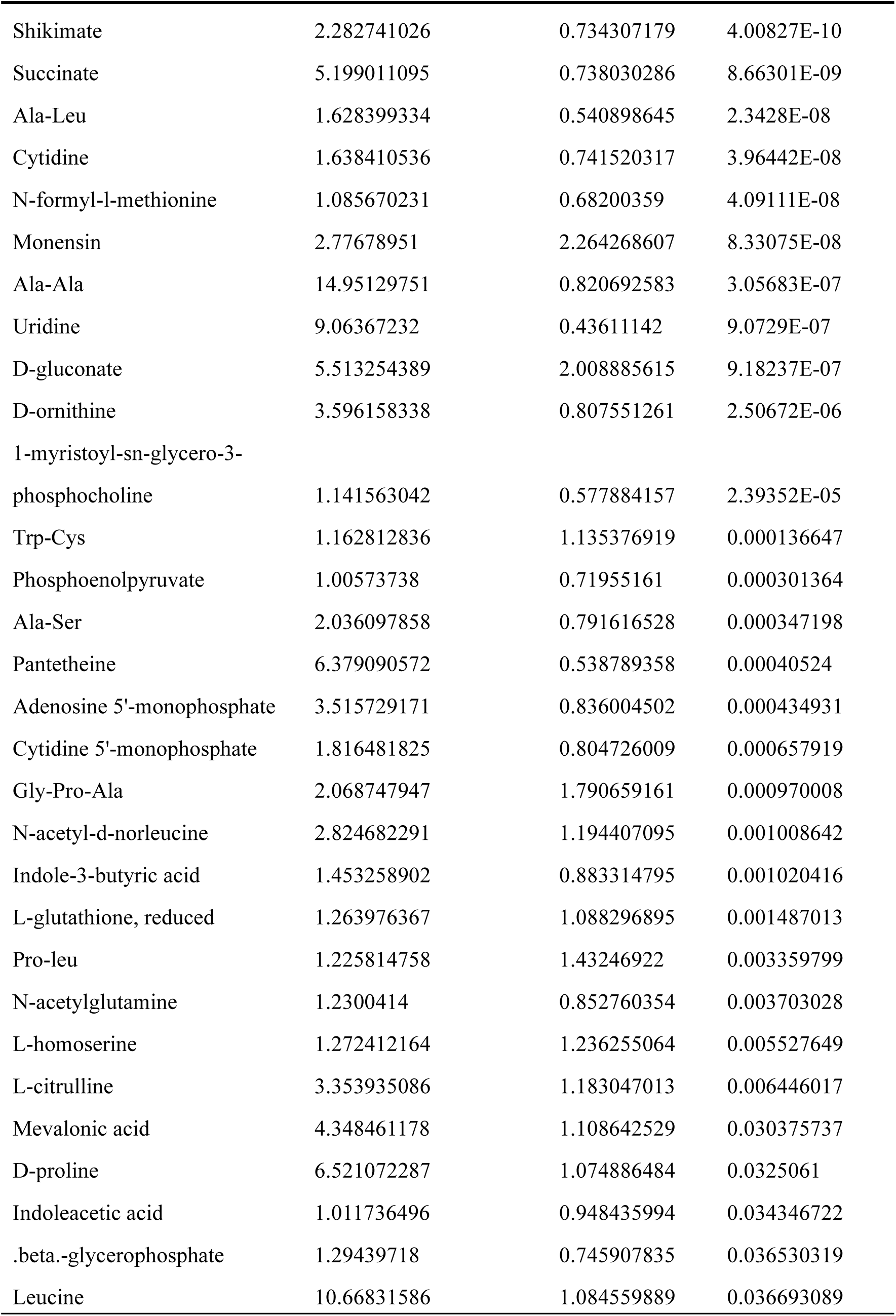
Staphylococcus aureus H7 vs H8 differential metabolites before freezing in negative ion mode.

**Table S3.**
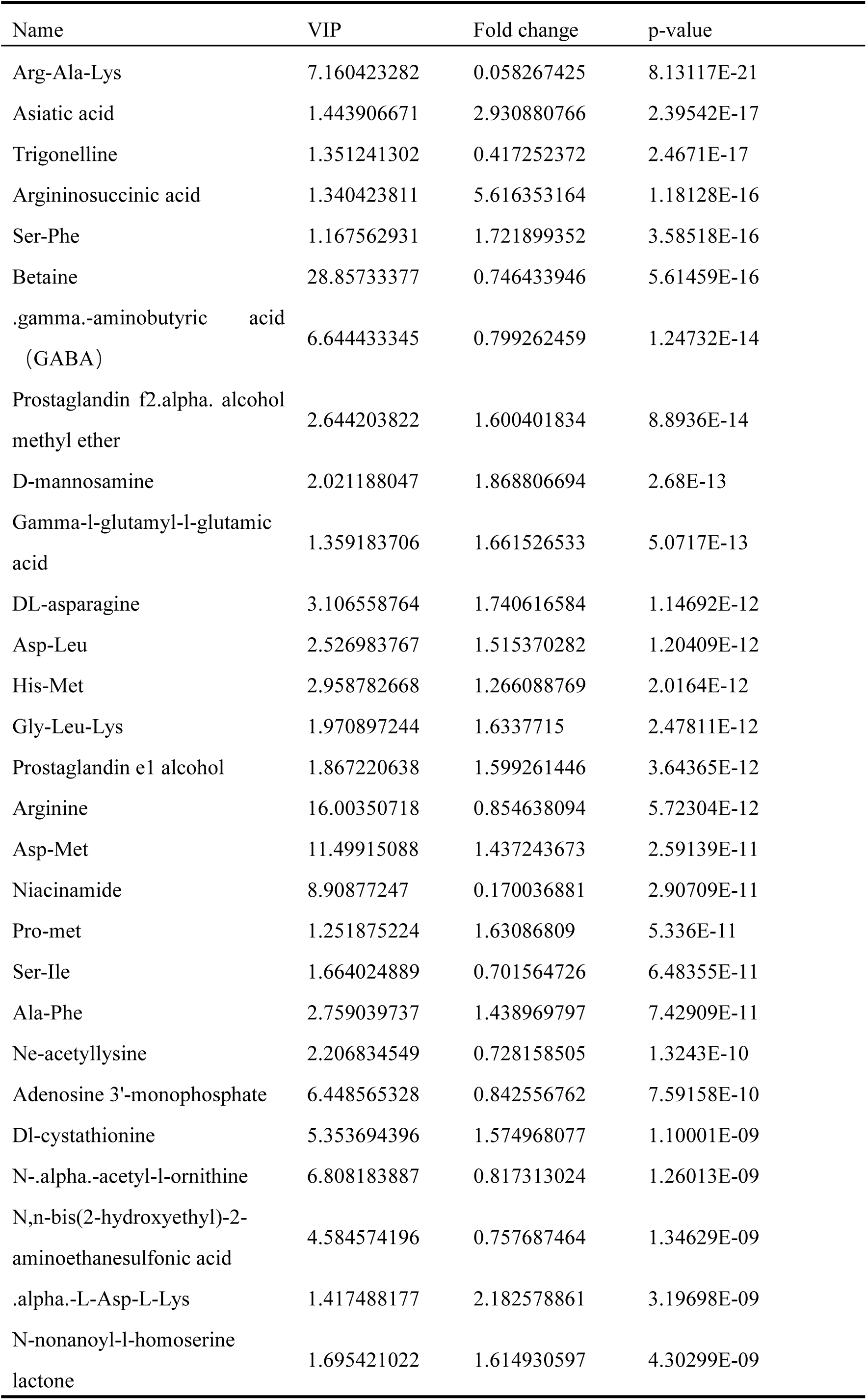
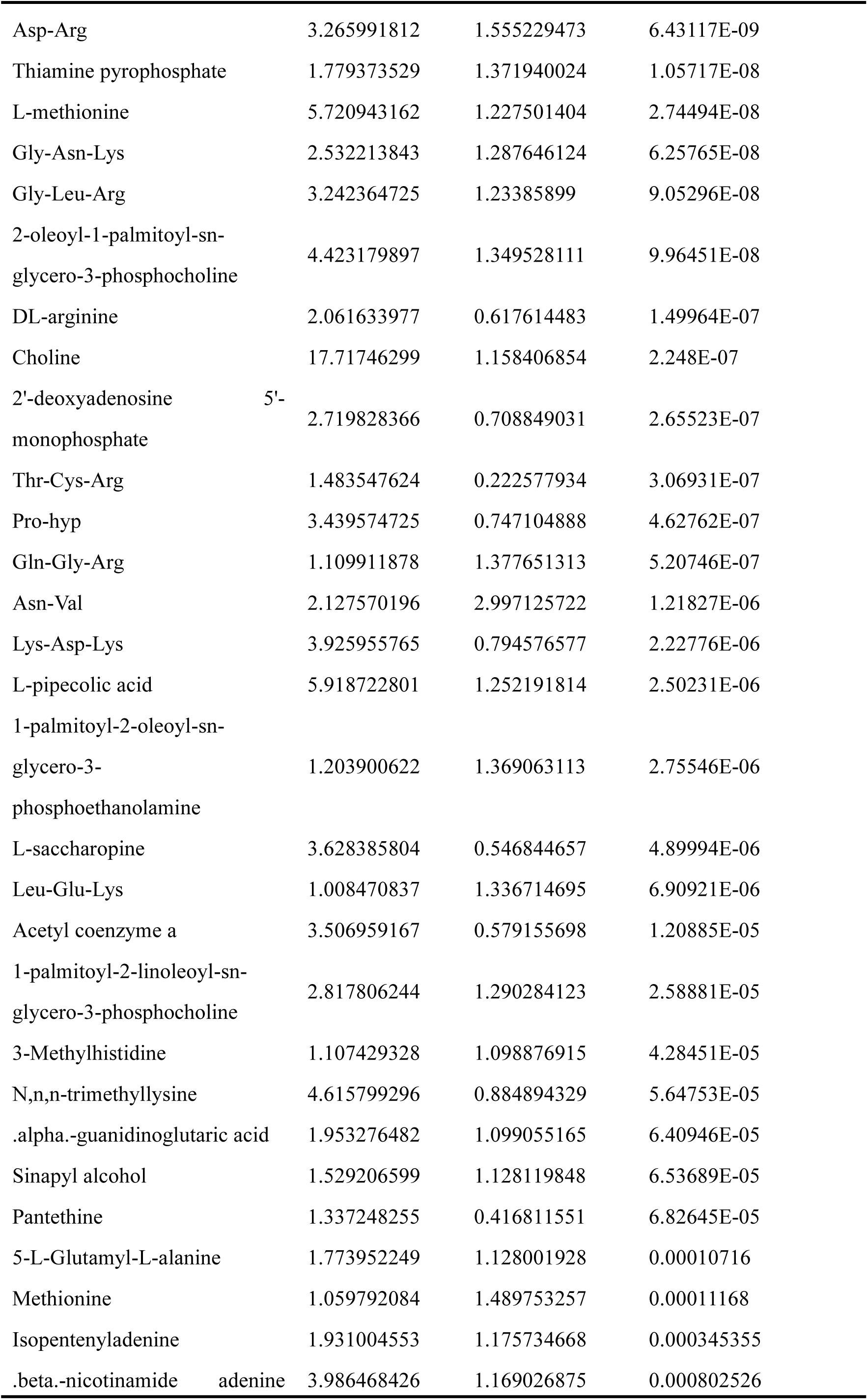
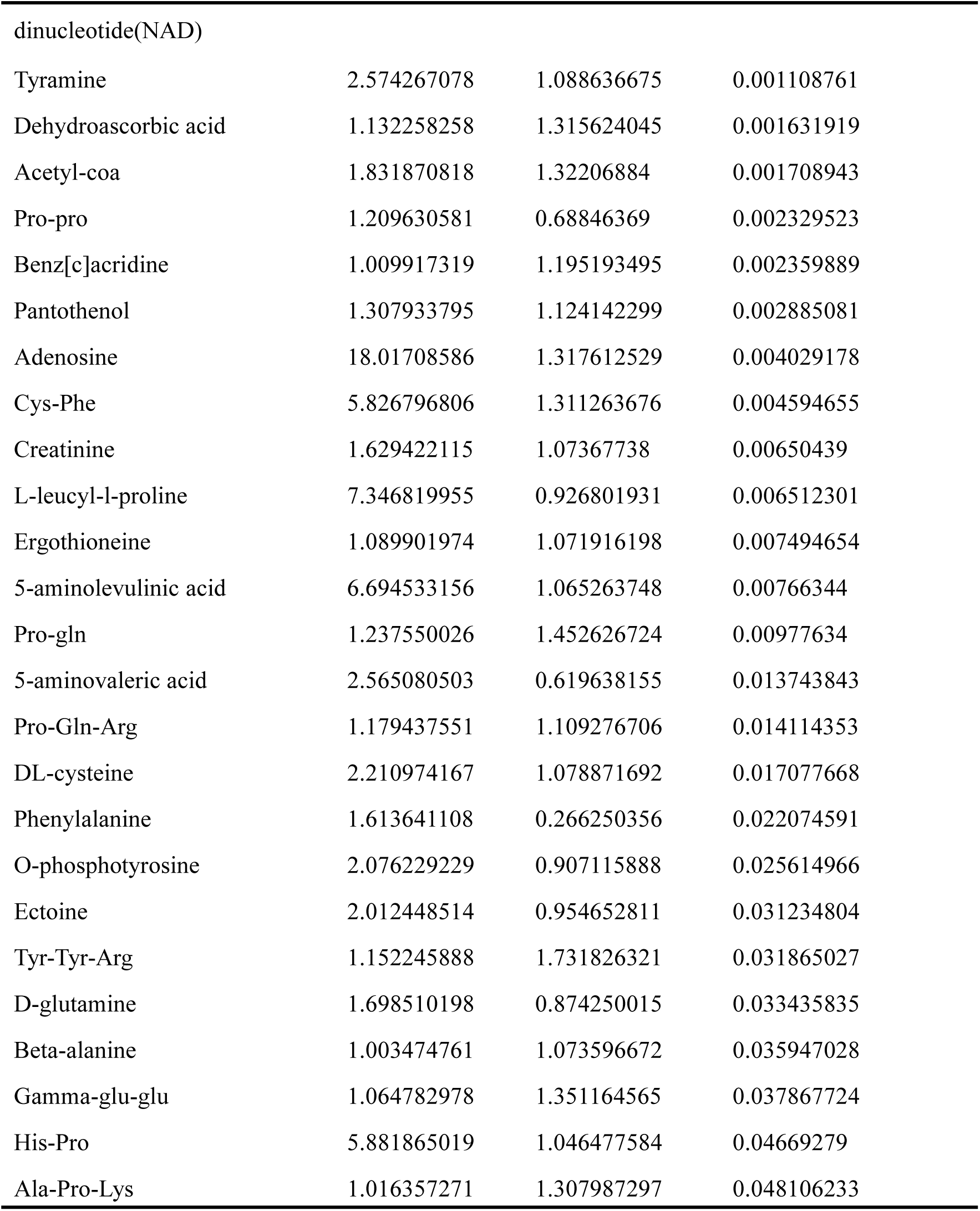
*Staphylococcus aureus* Q7 vs Q8 differential metabolites after freezing in positive ion mode.

**Table S4.**
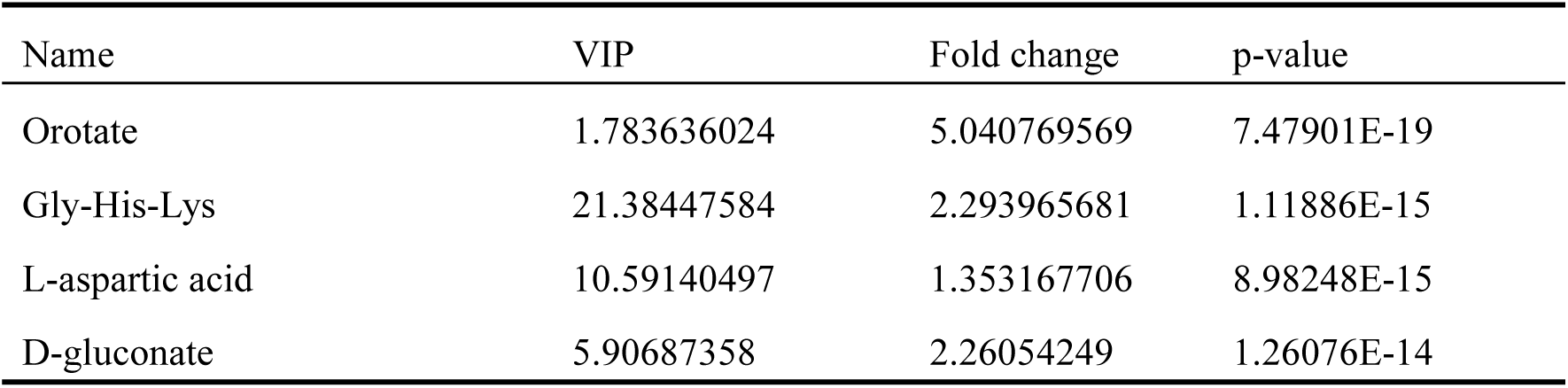
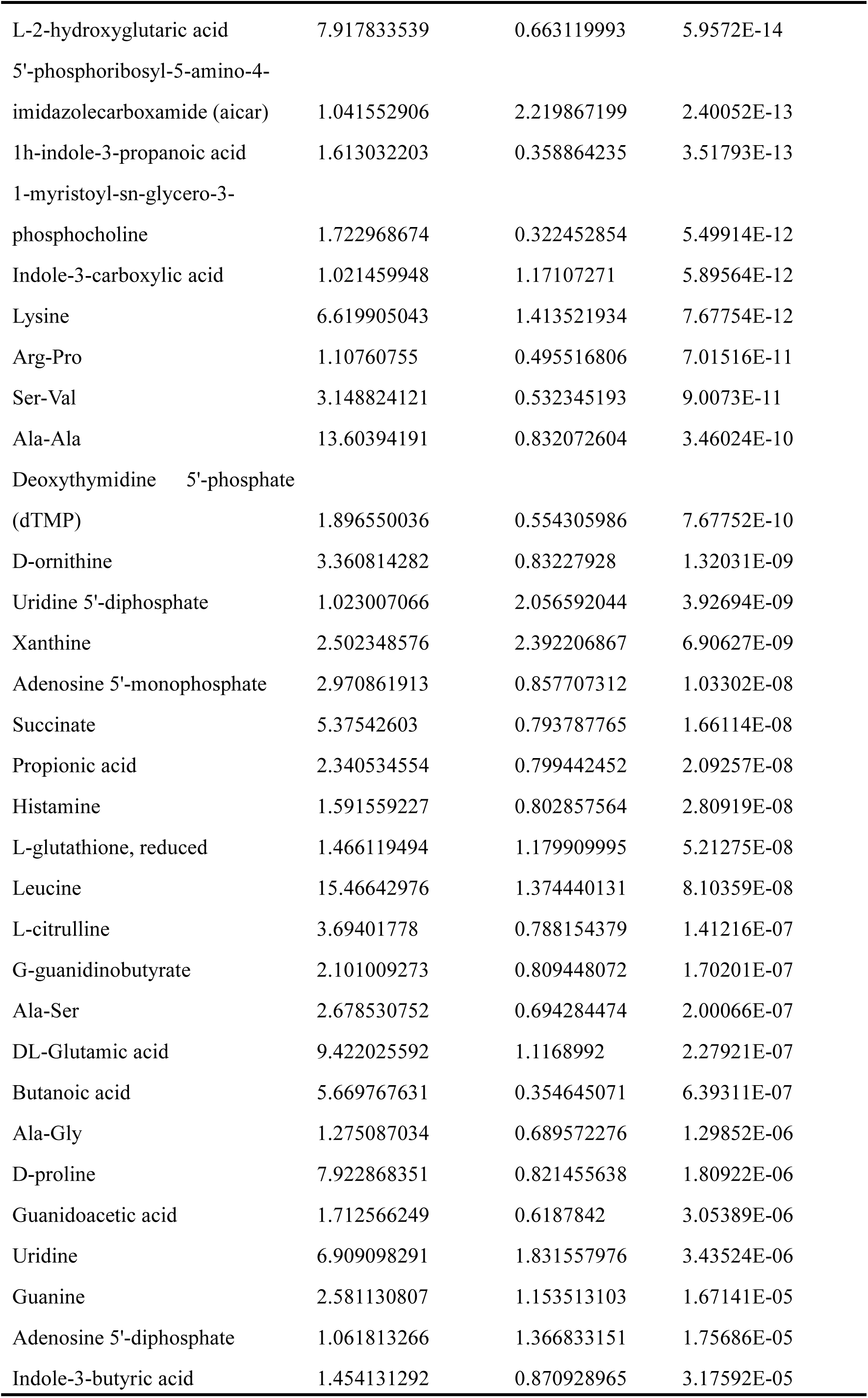
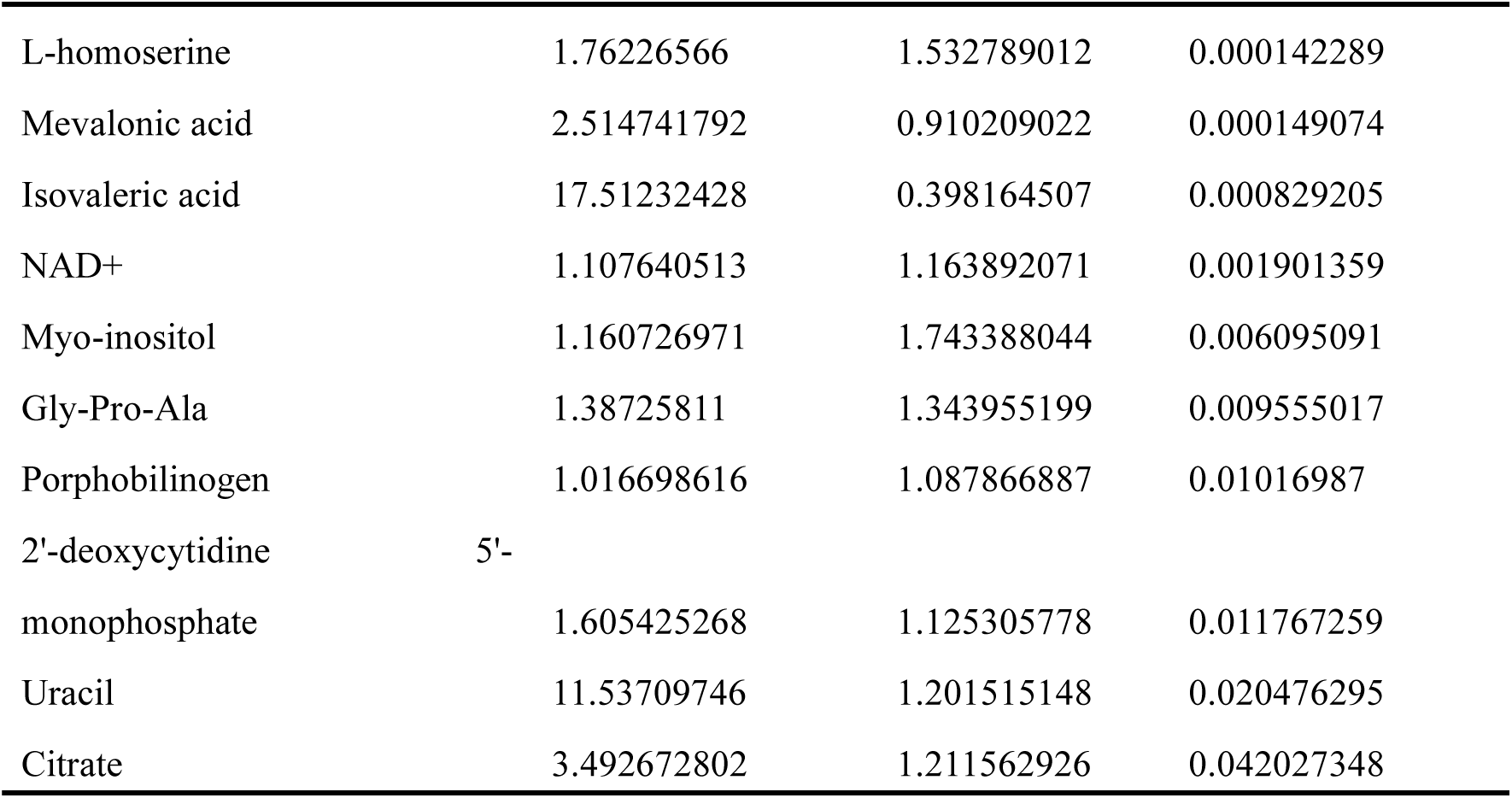
Staphylococcus aureus Q7 vs Q8 differential metabolites after freezing in nagetive ion mode.

